# RefSoil+: A reference for antimicrobial resistance genes on soil plasmids

**DOI:** 10.1101/435107

**Authors:** TK Dunivin, J Choi, AC Howe, A Shade

## Abstract

Plasmids harbor transferable genes that contribute to the functional repertoire of microbial communities, yet their contributions to metagenomes are often overlooked. Environmental plasmids have the potential to spread antibiotic resistance to clinical microbial strains. In soils, high microbiome diversity and high variability in plasmid characteristics present a challenge for studying plasmids. To improve understanding of soil plasmids, we present RefSoil+, a database containing plasmid sequences from 922 soil microorganisms. Soil plasmids were relatively larger than other described plasmids, which is a trait associated with plasmid mobility. There was no relationship between chromosome size and plasmid size or number, suggesting that these genomic traits are independent in soil. Soil-associated plasmids, but not chromosomes, had fewer antibiotic resistance genes than other microorganisms. These data suggest that soils may offer limited opportunity for plasmid-mediated transfer of described antibiotic resistance genes. RefSoil+ can serve as a baseline for the diversity, composition, and host-associations of plasmid-borne functional genes in soil, a utility that will be enhanced as the database expands. Our study improves understanding of soil plasmids and provides a resource for assessing the dynamics of the genes that they carry, especially genes conferring antibiotic resistances.

**Importance:** Soil-associated plasmids have the potential to transfer antibiotic resistance genes from environmental to clinical microbial strains, which is a public health concern. A specific resource is needed to aggregate knowledge of soil plasmid characteristics so that the content, host-associations, and dynamics of antibiotic resistance genes can be assessed and then tracked between the environment and the clinic. Here, we present RefSoil+, a database of soil-associated plasmids. RefSoil+ presents a contemporary snapshot of antibiotic resistance genes in soil that can serve as a reference as novel plasmids and transferred antibiotic resistances are discovered. Our study broadens our understanding of plasmids in soil and provides a community resource for investigating clinic-environment dynamics of important plasmid-associated genes, including antibiotic resistance genes.

## Introduction

Soil is a unique and ancient environment that harbors immense microbial biodiversity. The soil microbiome has functional consequences for ecosystems, like supporting plant growth (1, 2) and mediating key biogeochemical transformations (3). It also serves as a reservoir of microbial functional genes of interest to human and animal welfare. Within microbial genomes, important functions can be encoded on both chromosomes and extrachromosomal mobile genetic elements such as plasmids. Plasmids can be laterally transferred among community members, both among and between phyla (4–6). This causes propagation of plasmid functional genes and allows for them to spread among divergent host strains. Within microbial communities, plasmids influence microbial diversification (7) and contribute to functional gene pools (4). Plasmids can alter the fitness of organisms in a community as they can be gained or lost by environmental organisms, which alters their functional gene content and can have consequences for their local competitiveness.

Antibiotic resistance genes (ARGs) provide a prime example of the importance that functional genes encoded on plasmids can have. ARGs can undergo plasmid-mediated horizontal gene transfer (8, 9). There is particular concern about the potential for spread of ARGs between environmental and clinically-relevant bacterial strains. Studies of ARGs in soil have shown overlap between environmental and clinical strains that suggests HGT (10–12). For example, plasmid-encoded quinolone resistance (*qnrA*) in clinical Enterobacteriaceae strains likely originated from the environmental strain *Shewanella algae* (11). The extent of the impact of environmental reservoirs of ARGs is unknown (13), but studies have shown evidence for predominantly vertical, rather than horizontal, transfer of these genes (14). Additionally, it is speculated that rates of transfer in bulk soil are low compared to environments with higher population densities such as the rhizosphere, phyllosphere, and gut microbiomes of soil organisms (15). In the case of antibiotic resistance, mobilization is a public health risk. Broadly, the ability of plasmids to rapidly move genes both between and among membership is linked to diversification in complex systems, especially soils (7).

Despite their ecological and functional relevance, plasmids are not well characterized in soil. Plasmids vary in copy number, host range, transfer potential, and genetic makeup (4, 16), making them difficult to assemble and characterize from complex soil metagenomes that contain tens of thousands of bacteria and archaea (17). To aid in the study of plasmid-mediated transfer of functional genes in soil, we establish a resource to compare genetic locations of functional genes in soil organisms. We extended the RefSoil database (18) of 922 soil microorganisms to include their plasmids. We used this database to test whether soil-associated plasmids are distinct from plasmids from a broad, general database of microorganisms, RefSeq (19). We focused our comparisons on the content, diversity, and location of ARGs on plasmids and chromosomes. We used hidden markov models to search for clinically and agriculturally relevant ARGs in the extended soil database, RefSoil+, and RefSeq. RefSoil+ provides insights into the range of plasmid sizes and their functional potential within soil microorganisms. RefSoil+ can be used to inform and test hypotheses about the traits, functional gene content, and spread of soil-associated plasmids and can serve as a reference for plasmid assembly from metagenomes.

## Results and discussion

### Plasmid characterization

RefSoil+ is a database of soil-associated plasmids as an extension of RefSoil, which includes taxonomic information, amino acid sequences, coding nucleotide sequences, and GenBank files for a curated set of 922 soil-associated organisms. A total of 927 plasmids were associated with RefSoil organisms, and 370 RefSoil organisms (40.1%) had at least one plasmid (**Figure 1A**). This is high compared to the proportion of non-eukaryotic plasmids in the general RefSeq database (20%). The mean number of plasmids per RefSoil organism was 1.01, but the number of plasmids per organism varied greatly (**Figure 1B**). For example, strain *Bacillus thuringiensis* serovar thuringiensis (RefSoil 738) had 14 plasmids, ranging from 6,880 to 328,151 bp. The abundance of plasmids found in RefSoil genomes highlights plasmids as an important component of soil microbiomes (7, 20).

**Figure 1.**
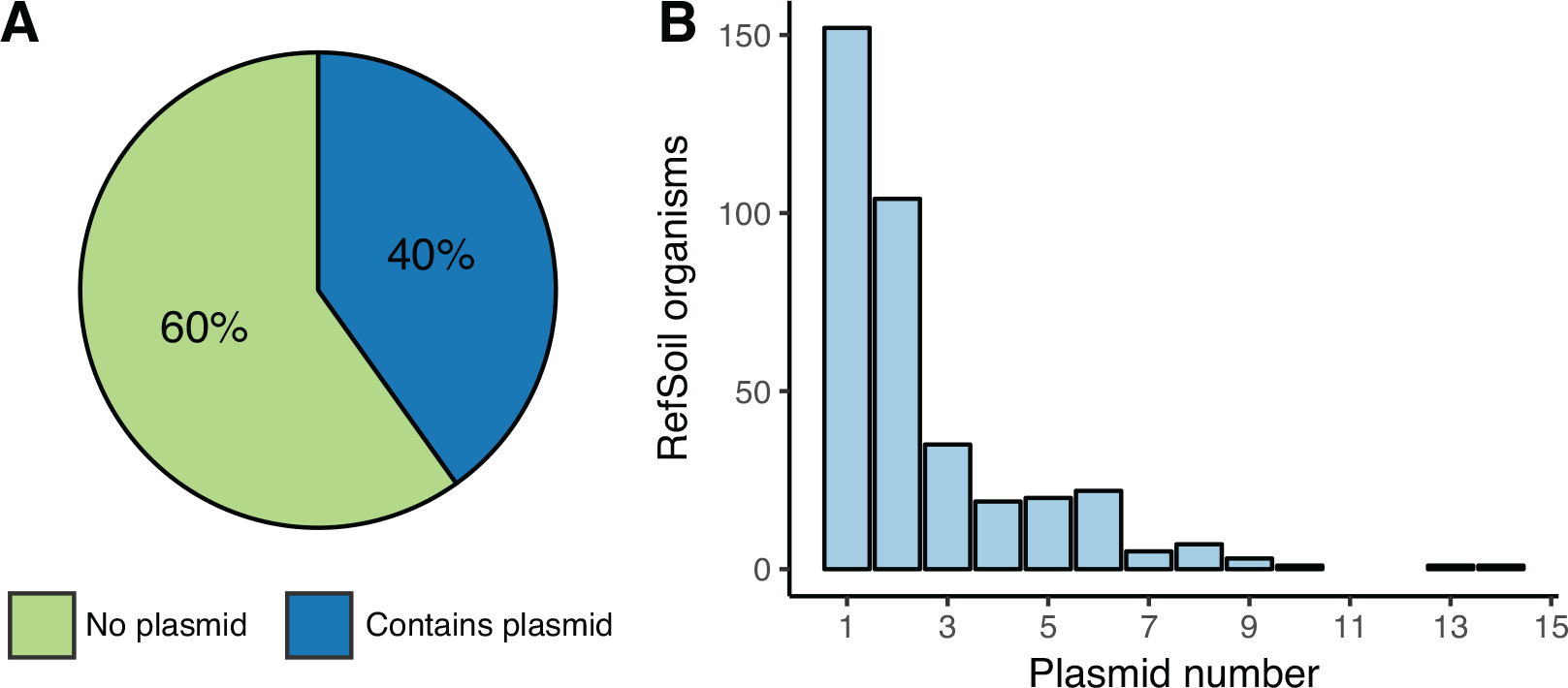
Summary of RefSoil plasmids. **A)**Percentage of RefSoil microorganisms with (blue) and without (green) detected plasmids. **B)**Distribution of the number of plasmids per RefSoil microorganism.

Soil-associated plasmids tended to be larger than plasmids from other environments. RefSoil plasmids contained > 195,000 kbp and increased the number of base pairs included in RefSoil by 4.4%. Plasmid size in RefSoil organisms ranged from 1,286 bp to 2.58 Mbp (**Figure 2A**), which rivals the range of all known plasmids from various environments (744 bp – 2.58 Mbp) (16). In the distribution of plasmid size, both upper and lower extremes had representatives from soil. Plasmids from all habitats had a characteristic bimodal size distribution with peaks at 5 kb and 35 kb (15–17). Soil-associated plasmids in RefSoil+, however, trended larger and did not have many representatives in the lower size range (**Figure 2**). Specifically, RefSoil+ proportionally contained more plasmids > 100 kb (**Figure 2B,** Mann-Whitney U test p < 0.001). Thus, while soil-associated plasmids vary in size, they are, on average, large. This is of particular importance because of the established differences in mobility of plasmids in different size ranges (5). Mobilizable plasmids, which have relaxases, tend to be larger than non-transmissible plasmids, with median values of 35 and 11 kbp respectively (5). The majority of soil-associated plasmids were > 35 kbp (**Figure 2**), suggesting they are more likely to be mobile. Additionally, conjugative plasmids, which encode type IV coupling proteins, have a larger median size (181 kbp) (5). The median size of soil-associated plasmids was 91 kbp (**Figure 2**), suggesting that these soil-associated plasmids are more likely to be conjugative. Future works should examine genetic potential for transfer of plasmids associated with different ecosystems to test this hypothesis.

**Figure 2.**
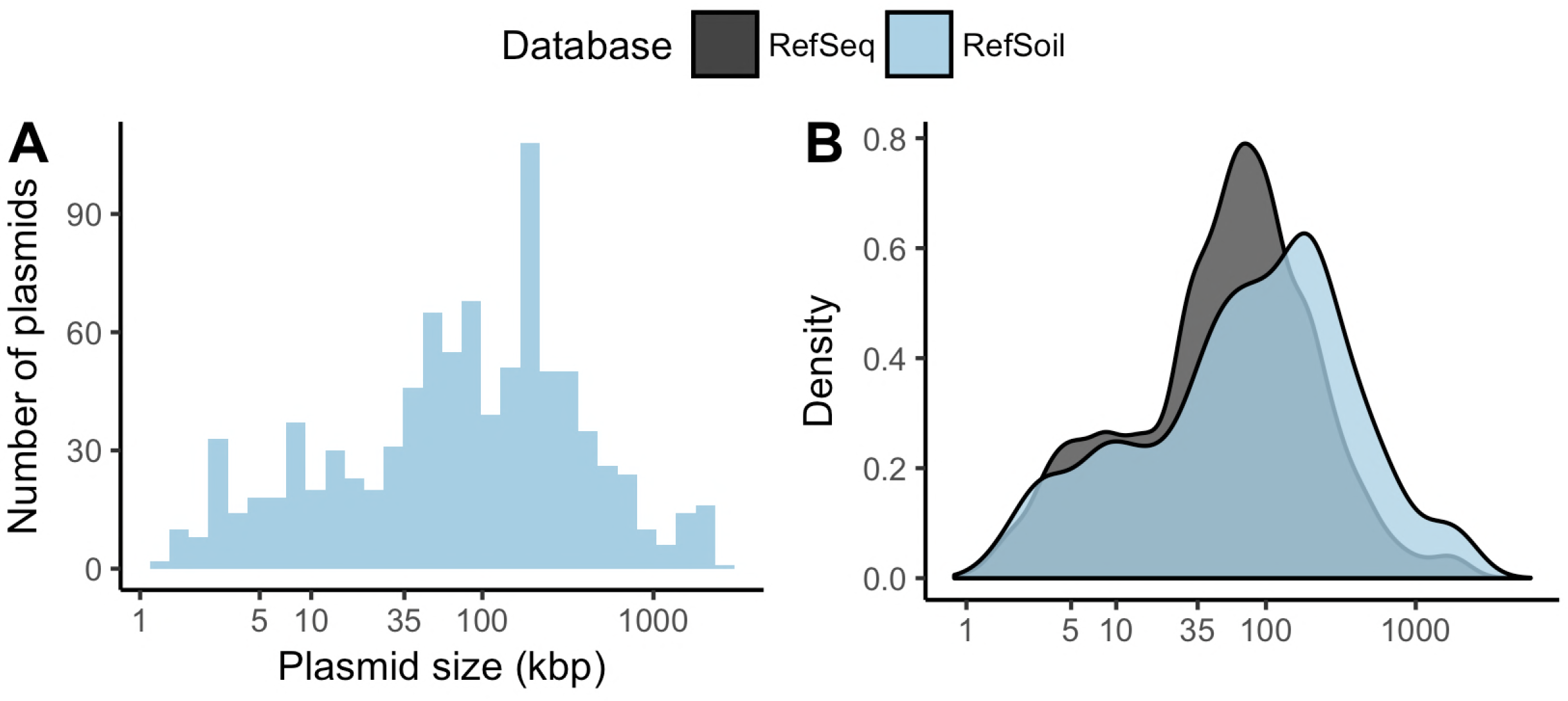
Plasmid size distributions. **A)**Histogram of plasmid size (kbp) from RefSoil plasmids. **B)**RefSoil (blue) and RefSeq (gray) plasmid size distributions.

Genome size, inclusive of chromosomes and plasmids, is an important ecological trait that is difficult to estimate from metagenomes (24). Due to incomplete assemblies, genome size must be approximated based on the estimated number of organisms through single-copy gene abundance (25). Extrachromosomal elements, however, inflate these estimated genome sizes because they contribute to the sequence information of the metagenome often without contributing single-copy genes (26). While our methodologies do not account for plasmid copy number (27), we examined the relationship between genome size and plasmid size in soil-associated organisms, and found none (**Figure 3**). Additionally, chromosome size was not predictive of the number of plasmids (**Figure 3**; **Figure S1**). For example, *Bacillus thuringiensis* subsp. thuringiensis Strain IS5056 had the most plasmids in RefSoil+, but these plasmids spanned the size range of 6.8 - 328 kbp. This strain’s plasmids make up 19% of its coding sequences (28), but its chromosome (5.4 Mbp) is average for soils (26). Despite that there is no clear relationship between genome size and plasmid characteristics within these data, the plasmid database can be used to inform estimates of average genome sizes from close relatives detected within metagenomes.

**Figure 3.**
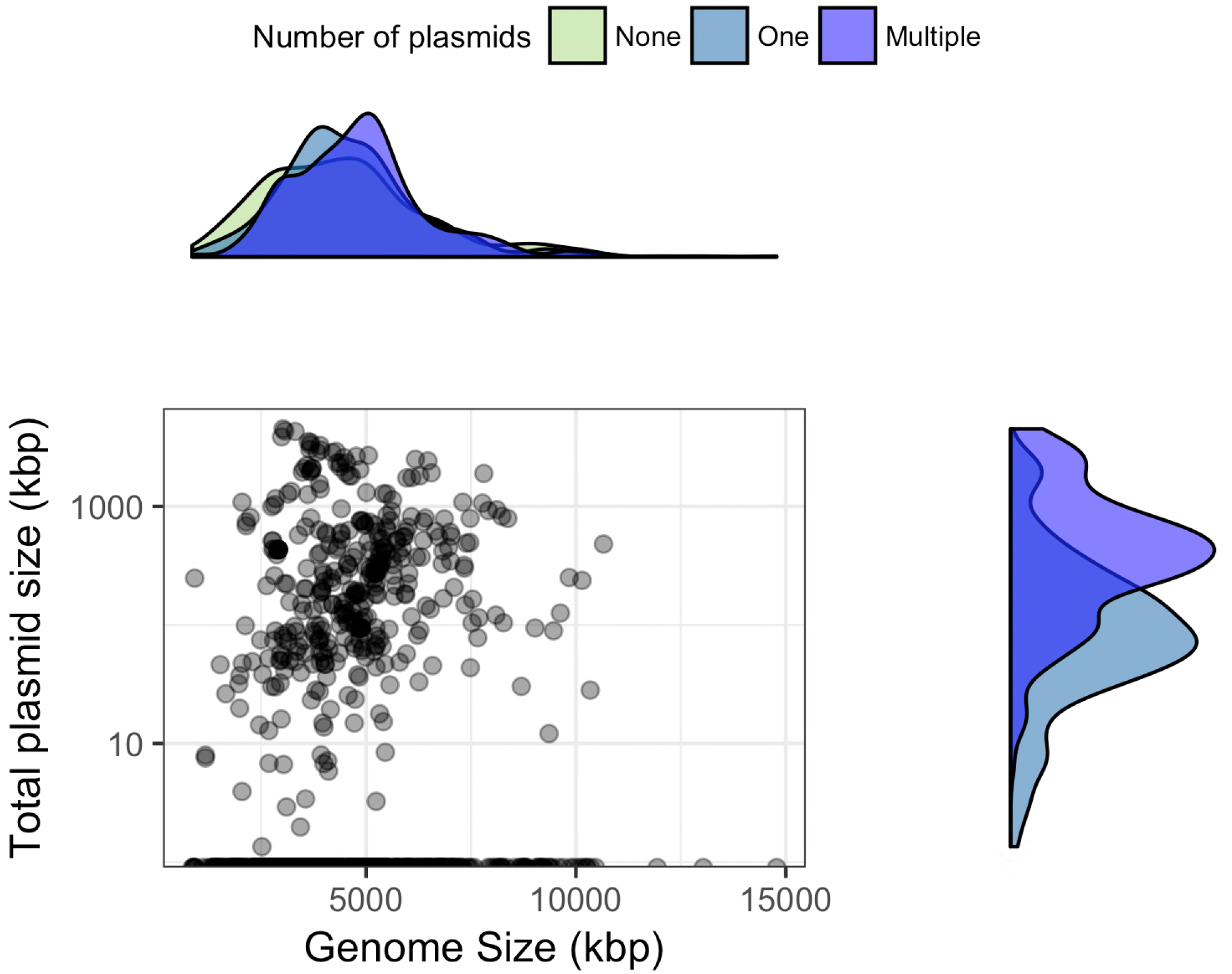
Relationship between plasmid size and genome size. Total plasmid size (sum of all plasmids in an microorganism, kbp) is plotted on a log scale against total genome size for each RefSoil microorganism. Density plots are included for each axis to represent the distribution of RefSoil microorganisms with different numbers of plasmids (none (green), one (blue), or multiple (purple)).

### ARGs in soil plasmids

It is unclear whether soil ARGs are predominantly on chromosomes or mobile genetic elements. While mobile gene pools are not static, there is evidence to suggest low transfer of ARGs in soil (14, 15, 29). For example, bulk soils are not a “hot spot” for HGT because they are often resource-limited (30), and surveys of ARGs in soil metagenomes have suggested a predominance of vertical transfer, rather than horizontal transfer, of ARGs (14, 29). Using RefSoil+, we examined 36 genes encoding resistance to beta-lactams, tetracyclines, aminoglycosides, chloramphenicol, vancomycin, sulfonamides, macrolides, and trimethoprim (29). After quality filtering, we detected 3,217 ARGs in RefSoil chromosomes and plasmids (**Figure 4; Table S1**).

**Figure 4.**
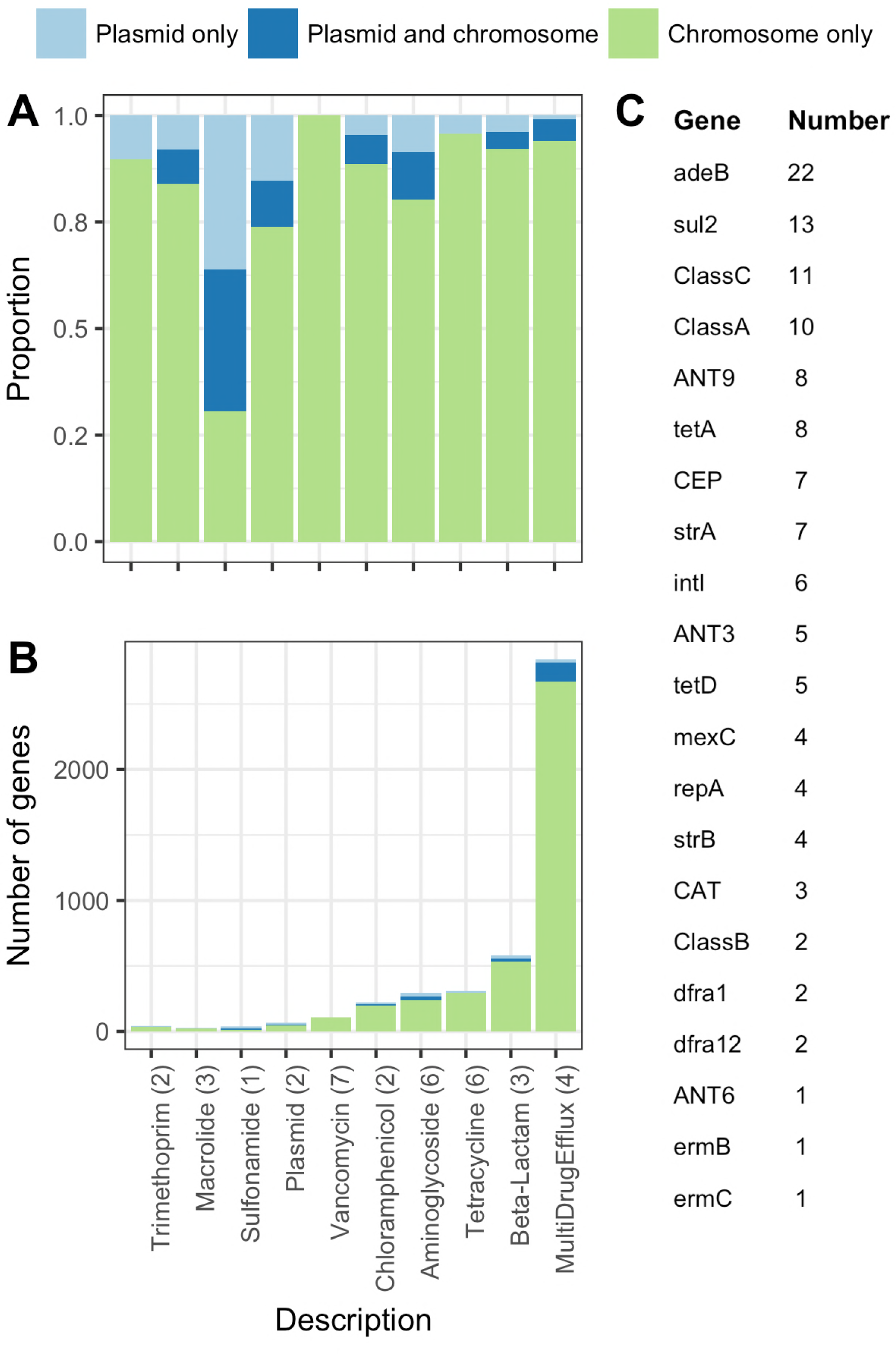
Distribution of ARGs in RefSoil genomes and plasmids. **A)**The proportion of ARGs on plasmids (light blue), genomes (green) or both (dark blue) in RefSoil+ microorganisms. **B)**The raw numbers of detected ARGs. Bars are colored by location of genomic element (as in panel A) and categorized by antibiotic resistance gene group. The number of genes included in each group is shown in parentheses. **C)**A table with the number different ARGs that were only found on plasmids. Genes are ordered by ranked abundance.

Adding plasmids to the RefSoil database increased functional genes in the database, as 128 ARG sequences were only detected on plasmids (**Figure 4C**). These functional genes would be missed if only chromosomes were considered. With the exception of sulfonamides, the majority of ARGs were chromosomally encoded in soils (**Figure 4AB**). We examined the genomic distributions of ARGs in RefSoil+ based on taxonomy (**Figure S3**). Proteobacteria had the most plasmid-associated ARGs, which has been reported previously (31). ARGs were found on chromosomes more often than plasmids, but we were curious whether this phenomenon was specific to soil. Therefore, we compared ARG content in RefSoil to all other known plasmids (RefSeq database; n = 9,132, (19)) and found that the number of ARGs per genome was comparable for RefSoil and RefSeq, but RefSoil plasmids had proportionally fewer ARGs than RefSeq plasmids (**Figure S4;**Mann-Whitney U test p-value = 0.002). This suggests that plasmid-mediated HGT rates of ARGs may be relatively low in these soil organisms. We note that the RefSoil database is limited in representatives of Verrucomicrobia and Acidobacteria which may change these estimates (18); however, this will improve as the database grows. We examined this trend for each gene individually and still observed a greater proportion of ARG sequences on plasmids in RefSeq compared with RefSoil+ with one exception, ANT9 which encodes a Streptomycin 3”-adenylyltransferase (**Figure 5**). Additionally, 12 genes (ANT3, CEP, *intI*, *qnr*, *repA*, *strA*, *strB*, *sul2*, *tetD*, *vanZ*) were more common on plasmids in RefSeq compared to only 3 genes (CEP, *dfra1*, *repA*) in RefSoil+ (**Figure 5**). Thus, these soil bacteria harbor relatively fewer ARGs on plasmids, suggesting that RefSoil+ organisms have limited capacity for plasmid-mediated transfer of these genes. These data represent a baseline of ARGs present on chromosomes and plasmids in soil microorganisms. This is important because some data suggest that soil ARGs are increasing over time due to increased antibiotic exposure (32). Future assessments of functional gene content on chromosomes and plasmids together will help to delineate changes in transfer potential and reveal selective or environmental factors that impact transfer potential.

**Figure 5.**
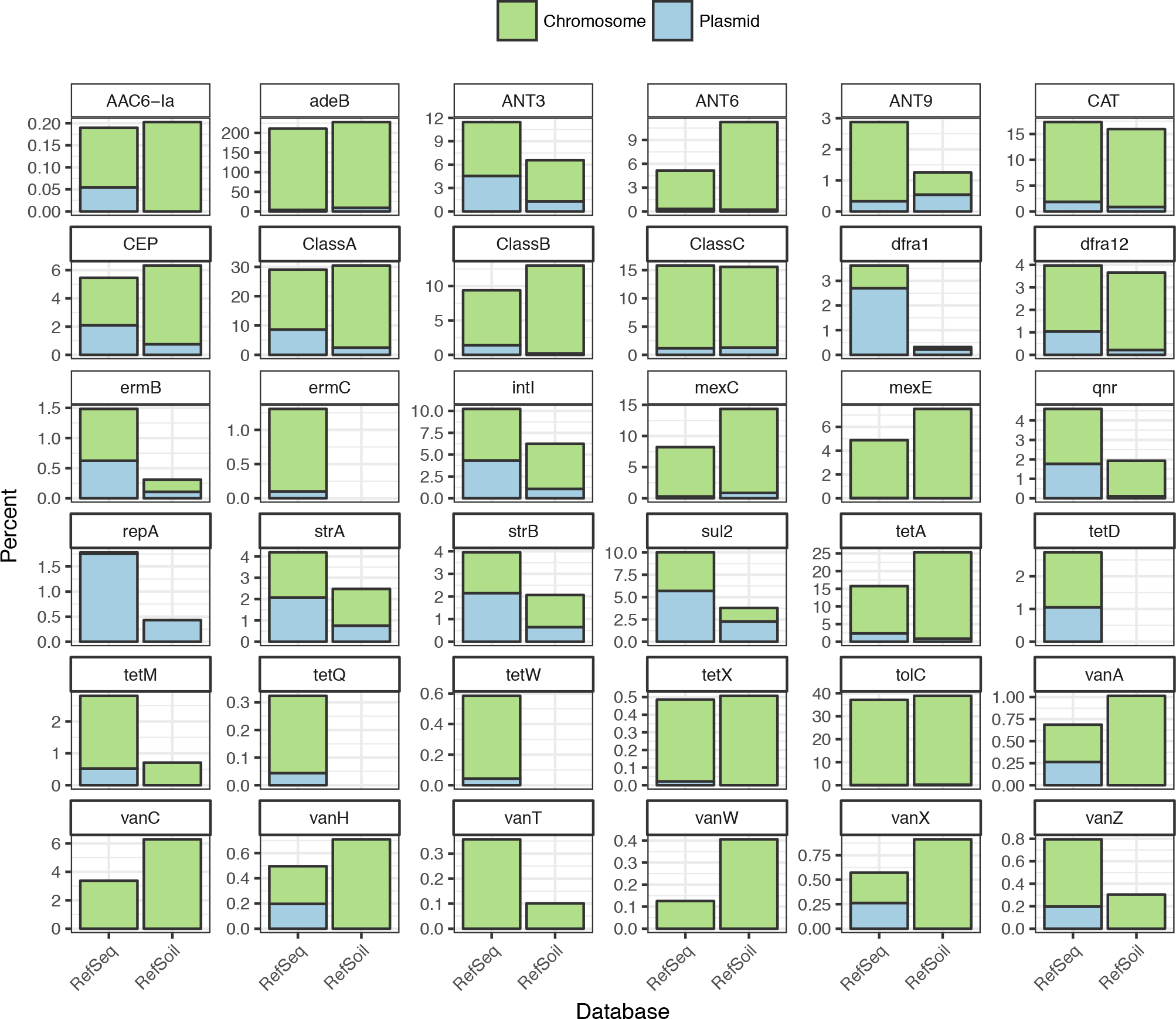
Proportion of genes on genomes and plasmids in RefSoil+ and RefSeq databases. Number of ARGs was normalized to number of genetic elements. Bars are colored by genetic element

We examined the abundance of ARGs in RefSoil+ and RefSeq strains and asked whether these ARGs were more commonly detected on chromosomes or plasmids. Gibson and colleagues (2015) compared soil-associated isolates with water and human-associated strains and found an abundance of genes encoding multidrug efflux pumps and beta lactam resistance but not tetracycline resistance in soil (33). This was also observed in our analysis (**Figure 5**). By determining whether ARGs were encoded on plasmids or chromosomes, we were also able to show that these patterns were due to chromosomal genes and more likely vertically transferred (**Figure 5**). While genome data from isolates cannot speak to environmental abundance of ARGs, our data support observations of ARGs in mobile genetic elements in soil from cultivation-independent studies as well. Luo and colleagues (2016) observed a low abundance of chloramphenicol, quinolone, and tetracycline resistance genes in soil mobile genetic elements (20), and Xiong and colleagues (2015) also observed low abundance of *qnr* genes in a soil mobile genetic elements (34). While plasmids are not the sole mobile genetic element, we observed fewer plasmid-encoded chloramphenicol, quinolone, and tetracycline resistance genes in soil-associated microorganisms than RefSeq microorganisms (**Figure 5**). Mobile genetic elements in soil have also been shown to have an abundance of genes encoding multidrug efflux pumps and resistance to beta-lactams, aminoglycosides, and glycopeptides (20). While we detected genes encoding aminoglycoside and beta-lactam resistance and multi drug efflux pumps in RefSoil+, we observed lower counts on plasmids as compared with chromosomes (**Figure 4**; **Figure 5**). Additionally, we did not detect plasmid-borne vancomycin resistance genes, despite that environmental samples have shown vancomycin resistance genes on mobile genetic elements (20). Though all isolate databases are biased by common cultivation conditions, these data point to gaps in our soil collections with a specific eye towards representation of plasmid content.

### RefSoil+ applications

Plasmid assembly tools rely on existing databases to assemble plasmids from metagenomes (35, 36), but this work shows that soil-associated plasmids are distinct. While this RefSoil+ is biased towards cultured strains, characterization of known plasmids is essential to improve detection of novel plasmids (21). This database of soil-associated plasmids expands knowledge of functional genes with potential for transfer in soil microbiomes, highlights the contribution of plasmids to metagenome-estimated genome size, offers insights into plasmid host ranges in soil, and serves as a reference for future works.

Host taxonomy can be observed in RefSoil+ because it is populated by the chromosomes and plasmids of isolates. While RefSoil+ does not predict plasmid presence or gene content in the environment, annotation of cultivable organisms with plasmids is important for soil systems because traditional methods of assembly and annotation from metagenomes allows only for coarse estimation of host identity (35, 37). Plasmid gene content is not static (38), and organisms can gain or lose plasmids (39, 40). Despite this, historical data of the genetic makeup and host range of plasmids can be used to better understand plasmid ecology, and to serve as an important reference to understand by how much host plasmid numbers and contents changes in the future.

RefSoil+ can be used to better target plasmids in the environment, whether it is used as a reference database or as a database for primer design. New microbiome sequencing techniques such as Hi-C sequencing (41), long-read technology (42), or single cell sequencing (43) could add to and leverage RefSoil+ to improve characterization of plasmid-host relationships in soil. As movement of ARGs are observed in the clinic and the environment, RefSoil+ can also serve as a reference for comparison with legacy plasmid and chromosome content and distributions. Novel genomes and plasmids could be added in future RefSoil+ versions, and plasmid-host relationships as well as encoded functions could be compared between cultivation-dependent and -independent methodologies. RefSoil+ provides a resource for research frontiers in plasmid ecology and evolution within wild microbiomes.

## Materials and methods

### Data availability

All data and workflows are publicly available on GitHub (http://github.com/ShadeLab/RefSoil_plasmids). A table of all RefSoil organisms with genome and plasmid accession numbers is available in **Table S2**and GitHub in the DATABASE_plasmids repository. This repository also hosts amino acid and nucleotide sequences for RefSoil+ genomes and plasmids. Plasmid retrieval workflows are included in the BIN_retrieve_plasmids directory. All workflows are included on Github as well in the ANALYSIS_antibiotic_resistance repository.

### RefSoil plasmid database generation

Accession numbers from RefSoil genomes were used to collect assembly accession numbers for all 922 strains. Assembly accession numbers were then used to obtain a list of all genetic elements from the assembly of one strain. Plasmid accession numbers were compiled for each strain and added to the RefSoil database to make RefSoil+ (**Table S1**). Plasmid accession numbers were used to download amino acid sequences, coding nucleotide sequences, and GenBank files. To ease comparisons between genome and plasmid sequence information, sequence descriptors for plasmid protein sequences were adjusted to mirror the format used for bacterial and archaeal RefSoil files.

### Accessing RefSeq genomes and plasmids

Complete RefSeq genomes and plasmids were downloaded from NCBI to compare with RefSoil. All RefSeq bacteria and archaea protein sequences were downloaded from release 89 (ftp://ftp.ncbi.nlm.nih.gov/refseq/release). All GenBank files for complete RefSeq assemblies were downloaded from NCBI. A total of 10,270 bacterial and 259 archaeal assemblies were downloaded. GenBank files were used to extract plasmid size and to compile a list of chromosomal and plasmid accession numbers. GenBank information was read into R and accession numbers for plasmids and chromosomes were separated. Additionally, all RefSoil accession numbers were removed from the RefSeq accession numbers. Ultimately, 10,359 chromosome and 9,132 plasmid accession numbers were collected to represent non-RefSoil plasmids. Protein files were downloaded and tidied using the protocol for RefSoil plasmids as described above.

### Plasmid characterization

We summarized the RefSoil+ and RefSeq plasmids in several ways. Plasmid size was extracted from GenBank files for each RefSoil genome and plasmid. For comparison, size was also extracted from RefSeq plasmids. These data were compiled and analyzed in the R statistical environment for computing (44). The RefSoil metadata (**Table S1**), which contains host information for each plasmid, was used to calculate proportions of RefSoil organisms with plasmids. Both the number of plasmids per organism and the number of RefSoil organisms with one plasmid were examined.

### Antibiotic resistance gene detection

We examined 36 clinically-relevant ARGs in RefSoil+, including *AAC6-Ia, adeB, ANT3, ANT6, ANT9, blaA, blaB, blaC, CAT, cmlA, dfra1, dfra12, ermB, ermC, intI, mexC, mexE, qnr, repA, strA, strB, sul2, tetA, tetD, tetM, tetQ, tetW, tetX, tolC, vanA, vanC, vanH, vanT, vanW, vanX,* and *vanZ*. For each gene of interest, hidden makrov models were downloaded from the FunGene database (45), which includes some models from the Resfams database (33). We then used these models to search amino acid sequence data from RefSoil genomes and plasmids with a publicly available, custom script and HMMER (46). To perform the search, hmmsearch (46) was used with an e-value cutoff of 10^−10^. These steps were repeated for protein sequence data from the complete RefSeq database (accessed 24 July 2018). Tabular outputs from both datasets were analyzed in R. Quality scores and percent alignments were plotted to determine quality cutoff values for each gene (**Figure S2**). All final hits were required to be within 10% of the model length and to have a score of at least 40% of the maximum score for that gene. Based on quality distributions and GenBank function assignments, additional quality filtering by score was applied to genes *adeB*, *CEP*, *vanA*, *vanC*, *vanH*, *vanX*, and *vanW*. When one amino acid sequence was annotated twice (i.e. for similar genes), the hit with the lower score was discarded. The final, quality filtered hits were used to plot the distribution of ARGs in RefSoil genomes and plasmids.

## Acknowledgements

AS acknowledges support in part from the National Science Foundation under Grants DEB #1655425 and DEB#1749544, from the USDA National Institute of Food and Agriculture and Michigan State AgBioResearch, and from the Great Lakes Bioenergy Research Center U.S. Department of Energy, Office of Science, Office of Biological and Environmental Research under Award number DE-SC0018409. TKD acknowledges support from the Michigan State University Department of Microbiology and Molecular Genetics Russell B. DuVall Fellowship. We thank the Jim Cole and the Ribosomal Database Project for helpful feedback on the work.

---

**Figure S1. Relationship between plasmid number and genome size.** Boxplots showing the distribution of genome sizes based on the number of plasmids. Numbers above boxplots show the number of organisms in that category. P-value from an ANOVA is also shown.

**Figure S2. Quality of RefSoil+ ARG hits.** Percent alignment was plotted against the score for each ARG hit for quality filtering purposes.

**Figure S3. Distribution of ARGs in RefSoil chromosomes and plasmids by taxonomy.** The number of detected ARGs were normalized to the number of RefSoil organisms in each phylum and Proteobacteria class. ARG hits are colored by genetic location. The number of taxa included in each phylum is shown in parentheses.

**Figure S4. Proportion of ARGs in RefSoil and RefSeq databases.** Boxplots of the proportion of ARGs per genetic element. Each ARG was normalized to the number of genetic elements in the database. Points are colored by ARG category, and P-values for Mann-Whitney U test are 0.55 (n.s. is not significant) and 0.007 (**) for chromosomes and plasmids respectively.

**Table S1.** Quality filtered ARG hits in RefSoil genomes and plasmids. Information on quality scores and accession numbers for each ARG hit.

**Table S2.** RefSoil taxonomy table with plasmid and genome accession numbers.

